# Layer-Wise Emergence of Human-Derived Object Dimensions in Deep Neural Networks

**DOI:** 10.1101/2025.05.19.654977

**Authors:** Florian Burger, Manuel Varlet, Genevieve L. Quek, Tijl Grootswagers

**Author notes:** **Corresponding Author:** Florian Burger.

## Abstract

Object recognition in the human visual system is implemented within a hierarchy characterised by increasing feature complexity. Here, we investigated whether human-derived dimensions of object knowledge become increasingly predictable from layer-wise activations in deep neural networks (DNNs), and how this predictability is related to architecture, learning objective, dimension type, and stimulus statistics. To test this, we predicted human-derived dimensions from layer-wise activations of multiple DNNs and transformer models trained on large-scale datasets. In trained DNNs, information relevant to human-derived object dimensions was largely absent in early layers, strengthened across layers, and peaked in later layers, in contrast to the flatter, non-progressive profiles we observed for raw pixel statistics and untrained networks. Differences in architecture, including recurrence and skip connections, and learning objectives were associated with changes in the strength and distribution of dimension-relevant information across layers. Dimension type also modulated predictability: perceptual dimensions were more predictable than semantic dimensions in simpler representations, a gap that was reduced or absent in more complex models. Stimulus statistics further modulated where this information was most available, including in a separate material perception dataset. These findings indicate that the layer-wise increase in the predictability of human-derived dimensions depends on training rather than on network depth alone, and that where in the network this information becomes available differs systematically with architecture, learning objective, dimension type, and stimulus statistics.

**Author Summary:** When we recognise an everyday object, our brain represents many properties about it, from simple visual features like colour and shape to more abstract properties like whether it is alive or how it is used. These properties, called human-derived dimensions, are measured from how people judge the similarity of objects. Deep neural networks, computer vision models inspired by the brain, are increasingly used to study how such knowledge might be represented artificially. We asked whether information about human-derived dimensions becomes progressively easier to detect as images pass through the layers of these networks. This pattern only emerged once networks had been trained. A network’s architecture, the task it was trained on, whether a property was simple or abstract, and the images it was shown were all associated with differences in where this information became most available. These findings show that information about object properties relevant to human perception becomes increasingly predictable across layers of deep neural networks, with the precise pattern depending on network architecture and training.

## Introduction

Recognising visual objects rapidly from retinal input is critical for everyday behaviour. The human visual system achieves this through a spatial–temporal hierarchy: simple features are extracted early in both space and time, while complex, abstract features emerge comparatively later. Early visual cortex such as V1 encodes local edges and orientations, whereas higher-level areas such as inferotemporal cortex (IT) represent complex object properties and categories (DiCarlo et al., 2012; Hubel & Wiesel, 1962). Time-resolved EEG suggest that early visual responses to objects are strongly driven by low-level image characteristics such as texture, with information about these low-level visual features peaking roughly 50–100ms after stimulus onset. Conversely, information about higher-level object properties such as animacy arises later during stimulus processing, peaking at around 150–200ms (Carlson et al., 2013; Robinson et al., 2023). Both low-level and high-level properties are relevant for behaviour, with low-level properties supporting rapid perceptual discrimination and high-level properties guiding conceptual understanding and goal-directed behaviour (Groen et al., 2018; Thorat et al., 2019). These behavioural relevance patterns can be captured in human-derived dimensions: continuous, data-driven axes estimated by modelling large-scale behavioural judgments, such as similarity ratings or odd-one-out choices, so as to best explain which objects humans treat as similar to one another (Hebart et al., 2020; Schmidt et al., 2025). Critically, a human-derived dimension is not itself a category: each dimension (e.g., “animacy” or “red”) is a continuous axis on which every object receives a continuous number indicating how strongly it expresses that dimension, rather than a label an object either does or does not belong to. Dimensions collectively span from perceptual features such as “red” to conceptual features such as “animacy”, together providing a high-dimensional, interpretable description of representational content. In the brain, these human-derived dimensions emerge in a systematic progression from early low-level dimensions to later high-level dimensions, reflecting the gradual transformation of visual information along the hierarchy (Contier et al., 2024; Teichmann et al., 2023).

Deep neural networks (DNNs) process information through a series of layers, with early layers aligning with early stages of visual processing in humans and later layers aligning with later stages in humans (Cichy et al., 2016; Guclu & Van Gerven, 2015; Yamins et al., 2014). In terms of human-derived dimensions, DNNs show a measurable overlap with human behaviour: Mahner et al. (2025) decomposed DNN similarity structure into latent visual and semantic dimensions (broadly corresponding to our perceptual and semantic dimension types) that align with human judgments. Furthermore, DNN activations contain decodable information about specific human-derived dimensions (Hebart et al., 2020; Kaniuth et al., 2024). Our question is closely related to a broader literature comparing DNN representations to biological vision more generally. Hong et al. (2016) showed that explicit, linearly decodable information about a set of experimenter-defined, category-orthogonal object properties (e.g., position, pose, size) increases along both the ventral stream and the layers of a hierarchical model, using a logic closely related to our own layer-wise approach. Other work has benchmarked which architectures best predict neural responses in visual cortex (Schrimpf et al., 2018), examined how well such models generalise to novel stimulus distributions (Ren & Bashivan, 2024), and characterised how architecture and training data shape high-level visual representations in brains and machines (Conwell et al., 2024); these studies largely quantify overall representational similarity or predictive accuracy at a given stage of processing, rather than treating the layer-wise trajectory of individual properties as the object of interest. It remains unclear whether information predictive of these human-derived dimensions follows a systematic layer-wise profile that resembles broad temporal and spatial progressions observed in the human visual system.

In this study, we map emergence profiles of human-derived object dimensions across DNN layers. We define an *emergence profile* as the layer-wise trajectory of prediction performance for a given human-derived dimension. Thus, an emergence profile indexes when and where DNN activations contain information sufficient to predict a dimension, without assuming that the model organises its internal representational space along the same axes or geometry as humans. Within the broader literature, our approach is closest in logic to Hong et al. (2016): both treat the layer-wise trajectory of specific properties, rather than an aggregate similarity or predictivity score, as the central unit of analysis. We depart from their approach in what these properties are: rather than a set of properties chosen to be independent of object category, we study a large, data-driven set of dimensions estimated directly from human similarity judgments, spanning from perceptual to semantic content and including dimensions closely tied to category membership itself (e.g., animacy). Mapping these emergence profiles allows us to compare the layer-wise availability of dimension-relevant information in DNNs with theoretical expectations from human vision.

To further understand what might shape emergence profiles, we draw on four factors known to influence representational organisation in human vision: architecture, learning objective, dimension type, and stimulus statistics. Architectural mechanisms in the brain such as recurrence, feedback, and skip connections refine responses in higher visual areas and shape the progression of representations across visual processing stages (Kar et al., 2019; Schmid et al., 2010; Spoerer et al., 2020). Learning objective further modulates processing: neurons in the visual cortex can adjust their tuning depending on behavioural goals, reflecting the influence of top-down objectives (Bracci et al., 2017; Harel et al., 2014). Dimension type itself may also matter: perceptual and semantic dimensions show distinct temporal and hierarchical signatures in the human brain (Carlson et al., 2013; Contier et al., 2024; Robinson et al., 2023; Teichmann et al., 2023), raising the question of whether perceptual and semantic human-derived dimensions show similarly distinct emergence profiles in DNNs (Mahner et al., 2025). Finally, the statistics of object inputs also matter, with different object domains accentuating different human-derived dimensions of visual information, such that object-focused datasets emphasise higher-level properties while material- and texture-focused datasets highlight perceptual features (Konkle & Caramazza, 2013; Hebart et al., 2020; Schmidt et al., 2025). Yet how much each of these factors contributes to the layer-wise profile of dimension-relevant information remains unclear. DNNs provide a powerful testbed to examine these influences systematically (Kriegeskorte, 2015). By comparing emergence profiles across systematic manipulations of network architecture, learning objective, and stimulus statistics, and by directly contrasting perceptual and semantic dimensions, we can probe how these four factors shape emergence profiles. Although these manipulations are artificial, they provide a principled way to anticipate how similar factors might influence the emergence of human-derived dimensions in human vision.

In this study, we extracted layer activations from DNNs and used regression models to predict human-derived dimensions from these activations. This produced an emergence profile for each dimension, capturing how prediction performance changed across layers. We then examined how these emergence profiles are related to four factors: network architecture, learning objective, dimension type, and stimulus statistics. To anticipate our results, we found that emergence profiles depended on training: prediction from raw pixel statistics and untrained networks remained flat and weak across layers, whereas in trained networks, high-level dimensions such as animacy showed weak early-layer prediction, stronger mid-layer prediction, and peak prediction in later layers. Recurrence and skip connections were associated with greater magnitude and persistence of dimension prediction. Learning objective was associated with differences in the overall distribution of dimension-relevant information across layers. A predictability advantage for perceptual over semantic dimensions was present in simpler representations but diminished in more complex models. Finally, for a material-focused stimulus set, dimension-relevant information was shifted toward earlier layers relative to the object-dataset profile, while still peaking toward the top of the network. Our contribution therefore goes beyond showing that human-derived dimensions can be decoded from DNN activations: we show that the layer-wise increase in decodability depends on training rather than on depth alone, and we locate where in the network, and for which dimensions, this increase differs between models and stimuli. By treating emergence profiles as the central unit of analysis, we link what can be predicted from DNN activations with how this prediction varies across these four factors. This provides a layer-resolved account of when and where information relevant to human-derived dimensions is available within artificial systems.

## Results

To examine whether DNN activations contain information predictive of human-derived dimensions in a systematic layer-wise manner, we analysed representations across a range of deep neural networks, from feedforward models to recent transformer models. We used images from two datasets: the THINGS object dataset (Hebart et al., 2019), which contains a broad range of everyday objects, and the STUFF material perception dataset (Schmidt et al., 2025), which focuses on natural and man-made materials (Figure 5A). In comparison, THINGS emphasises high-level human-derived object dimensions such as animacy and tool-relatedness, whereas STUFF highlights perceptual human-derived material dimensions such as colour and texture (Hebart et al., 2020; Schmidt et al., 2025). For each model and dataset, we extracted activations from every layer, applied principal component analysis (PCA) at each layer to reduce dimensionality, and used ridge regression to predict the relevant human-derived dimensions from these components (Figure 1A, see Methods). All primary analyses reported below were also replicated under ridge, unconstrained linear, and nonnegative regression as well as under alternative PCA configurations (50 and 100 fixed components, 95% variance threshold), indicating that our conclusions do not depend on specific analytic choices (see Methods, S2 Fig).

**Figure 1.**
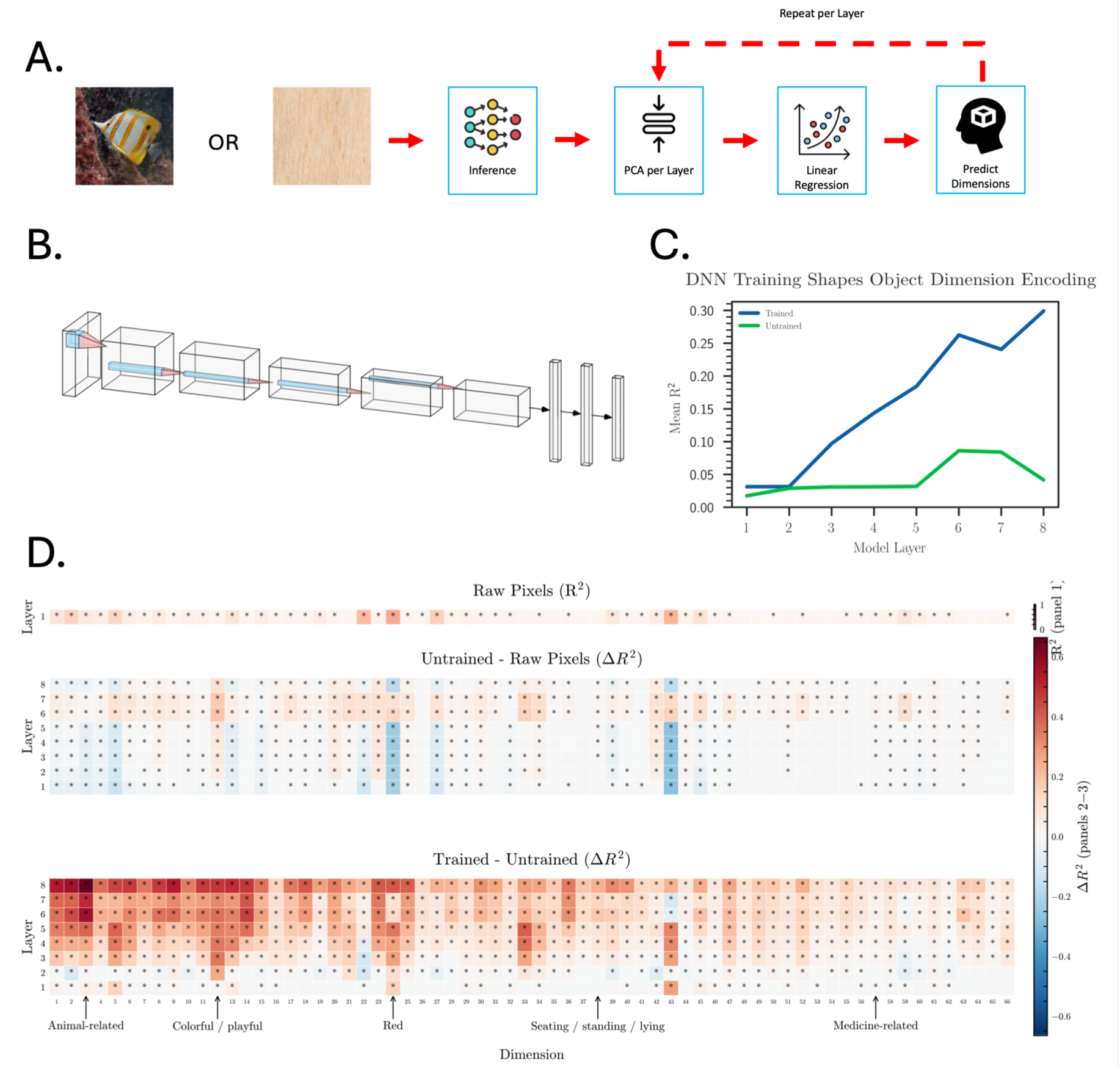
Predicting human-derived dimensions from images and DNN activations. **A.** Workflow illustration for both THINGS and STUFF datasets: images are passed through the network, PCA is applied to each layer, and ridge regression is used to predict human-derived dimensions (repeated per layer). Object image: THINGSplus (CC0; Stoinski et al., 2023). Material image: “Balsa Wood Texture 1,” Joseph Francis (CC BY 2.0). **B.** Schematic architecture of AlexNet. **C.** Mean prediction accuracy (R²) across layers for trained (blue) and untrained (green) AlexNet, showing that training induces a gradual increase in dimension predictability while untrained remains flat/consistent. **D.** Heatmaps showing prediction accuracy (R²) and model differences across human-derived object dimensions. Top row: baseline performance from raw pixel values. Middle row: difference between untrained model and raw pixels. Bottom row: difference between trained and untrained model. Colour indicates strength/difference of prediction accuracy (R^2^), with asterisks marking permutation-tested significance (FDR-corrected; see Methods).

To test whether layer-wise prediction profiles depend on learning, or whether similar profiles are already present in the input or induced by network architecture alone, we first compared three baseline cases: raw image pixels, an untrained DNN, and a trained DNN. This initial analysis tests whether a systematic layer-wise increase in dimension prediction can arise without learning, or whether training is required for DNN activations to produce emergence profiles with a layer-wise increase in prediction. As the DNN, we used AlexNet, a widely used architecture in cognitive science (Figure 1B; Krizhevsky et al., 2012). Prediction from raw pixels showed that some perceptual dimensions, such as “Red,” which captures variation in perceived redness across objects, are already reflected in stimulus statistics, whereas high-level dimensions such as “Seating / Standing / Lying-related” showed little to no prediction strength, indicating limited high-level structure (Figure 1C-D). The untrained model showed slightly higher prediction for some higher-level dimensions than raw pixels, consistent with weak architectural biases, but prediction remained dominated by perceptual features and did not show a clear layer-wise emergence profile (Figure 1C-D). In contrast, the trained model exhibited a clear layer-wise emergence profile: information predictive of human-derived object dimensions was weak or absent in the earliest layers, became more decodable in mid-layers, and peaked toward the top of the network (Figure 1C-D). Thus, unlike raw pixels or an untrained network, a trained DNN shows an emergence profile in which information predictive of human-derived dimensions becomes increasingly available across model depth.

### Architecture: Emergence Profiles Are Found Across All Models and Are Stronger with Brain-Like Mechanisms

Next, we investigated whether architectural choices influence emergence profiles, as different architectures alter how information flows through a network (Kar et al., 2019; Spoerer et al., 2020). To test this, we analysed three architectures from the CORnet family (Kubilius et al., 2018), which share an identical four-layer layout but differ in their inclusion of within-layer recurrence and within-layer skip connections (Figure 2A). Across all three architectures, information predictive of human-derived object dimensions followed a broadly similar layer-wise profile: prediction was weak or absent in early layers, increased in mid-layers, and peaked toward the top of the network (Figure 2B).

**Figure 2.**
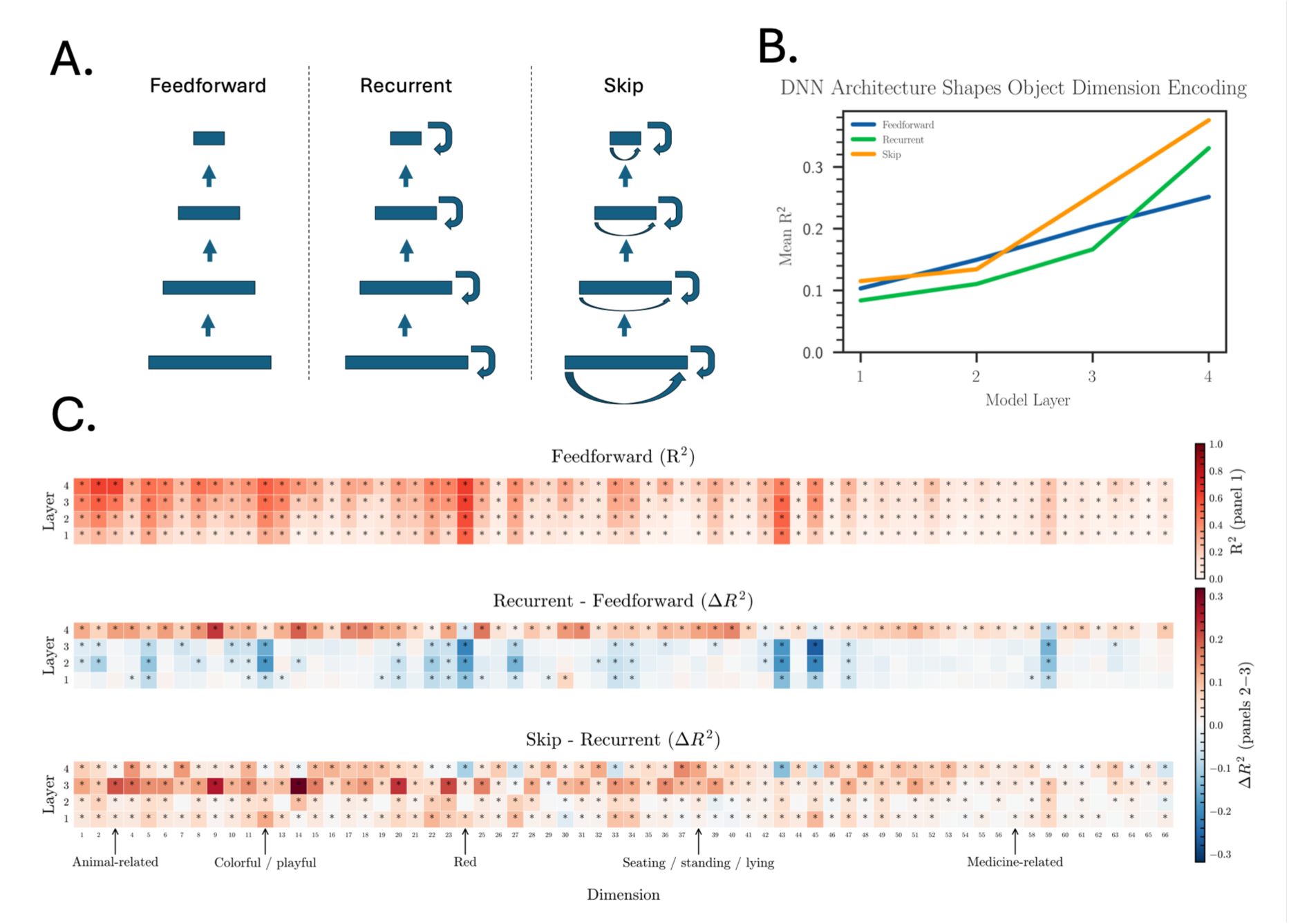
Comparing feedforward, recurrent, and skip architectures in predicting human-derived object dimensions. **A.** Schematic illustrations of three model types: feedforward (no recurrence or skip connections), recurrent (within-layer recurrence), and skip (within-layer skip connections). **B.** Mean prediction accuracy (R²) across model layers (V1–IT equivalent, see Kubilius et al. 2018) for each architecture, showing higher prediction of human-derived object dimensions in recurrent and skip networks relative to feedforward. **C.** Heatmaps showing prediction accuracy and architectural differences across the 66 dimensions. Top: baseline accuracy (R²) for the feedforward model. Middle: difference in prediction accuracy (ΔR²) between recurrent and feedforward architectures. Bottom: difference between skip and recurrent architectures. Colour indicates strength/difference of prediction accuracy (R^2^), with asterisks marking permutation-tested significance (FDR-corrected; see Methods).

Relative to the feedforward model (CORnet-Z), the recurrent model (CORnet-RT) showed stronger prediction of human-derived object dimensions in the final layer (Figure 2B–C). While the overall emergence profile shape was similar, recurrence was associated with a greater magnitude of dimension prediction, consistent with the possibility that recurrent processing makes dimension-relevant information more available or persistent across layers (Figure 2B–C). The third model, which combines recurrence with skip connections (CORnet-S), produced the strongest emergence profiles, with prediction performance rising earlier and reaching higher peak values than in the other models (Figure 2B–C). This pattern suggests that skip connections may facilitate earlier availability of dimension-relevant information across the hierarchy.

Together, these results show that information predictive of behaviourally relevant human-derived object dimensions is available across DNN architectures and follows broadly similar emergence profiles. Recurrence and skip connections were systematically associated with stronger profiles, suggesting that brain-inspired architectural mechanisms may modulate the layer-wise availability of dimension-relevant information.

### Learning Objective: Emergence Profiles Vary with Learning Objectives

Next, we examined how the emergence profiles of human-derived object dimensions from DNN activations differ across learning objectives, motivated by evidence that task demands modulate representational structure in biological vision and DNNs (Bracci et al., 2017; Zhuang et al., 2021). To do this, we compared models with similar transformer-based architectures but different learning objectives (Figure 3A): a vision transformer (ViT; Dosovitskiy et al., 2021), trained with a supervised visual classification objective; the vision branch of CLIP (Radford et al., 2021), trained with an image–text contrastive objective; and a self-supervised vision model (DINOv3; Siméoni et al., 2025).

**Figure 3.**
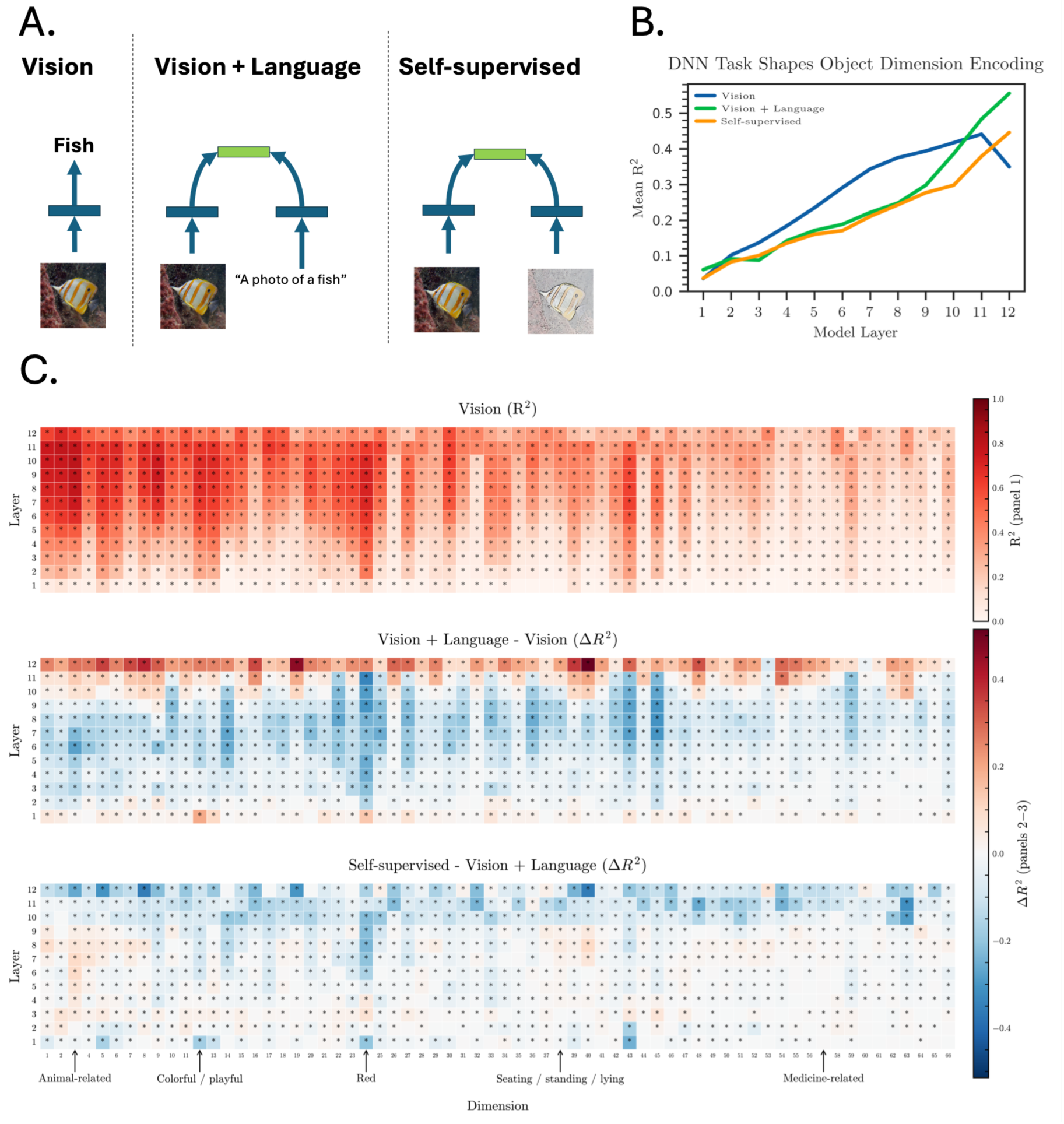
Comparing learning objectives (vision, vision–language, self-supervised) in predicting human-derived object dimensions. **A.** Schematic illustrations of three training paradigms: vision-only (standard supervised image classification), vision + language (image–text contrastive training, e.g., “a photo of a fish”), and self-supervised vision (self-supervised distillation). Object image: THINGSplus (CC0; Stoinski et al., 2023). **B.** Mean prediction accuracy (R²) across layers for the three models. CLIP (vision + language; green) shows higher prediction accuracy in later layers compared to supervised ViT (blue) and self-supervised DINOv3 (orange). **C.** Heatmaps showing prediction accuracy and differences between learning objectives across the 66 dimensions. Top: baseline accuracy (R²) for supervised ViT. Middle: difference in prediction accuracy (ΔR²) between CLIP and ViT. Bottom: difference between DINOv3 and CLIP. Colour indicates effect size, with asterisks marking permutation-tested significance (FDR-corrected; see Methods).

Across all three models, human-derived object dimensions were differentially predictable across layers (Figure 3B–C). In the vision-only model (ViT), emergence profiles increased across early and mid-layers but declined toward the final layer, consistent with later classification-specific layers becoming less informative for predicting the full set of human-derived dimensions (Figure 3B–C). In contrast, the vision–language model (CLIP) showed a steady increase in prediction performance across layers, peaking in the final layers that map visual features into a shared image–text embedding space (Figure 3B–C). The self-supervised model (DINOv3) followed a similar trajectory to the vision–language model, with strong increases across early and mid-layers with prediction performance peaking again in the final layer (Figure 3B–C).

Together, these results show that layer-wise prediction of human-derived object dimensions is observed across visual models, but that the shape and distribution of these emergence profiles differ with the model’s learning objective. Different learning objectives may emphasise different types of information, such as categorical, conceptual, or invariant information, which may produce systematic differences in when and where human-derived dimensions are predictable from DNN activations.

### Semantic vs. Perceptual: Differences Are Strongest in Simpler Representations

A key remaining question is whether the human-derived object dimensions can be divided into meaningful sub-groups based on whether they capture perceptual or semantic properties. Perceptual dimensions describe lower-level visual attributes of a stimulus, such as “red,” “white,” “circular/round,” or “long/thin,” whereas semantic dimensions describe conceptual or category-level properties, such as “animal-related,” “tool-relatedness,” “food-related,” or “electronics/technology.” We classified each of the 66 dimensions into perceptual or semantic groups following Mahner et al. (2025), yielding 15 perceptual and 44 semantic dimensions; the remaining 7 dimensions were of mixed or unclear type and were excluded from this analysis.

We then asked whether the emergence profiles of these two groups diverged across model layers. For each model, we compared mean prediction accuracy (R²) for perceptual and semantic dimensions at every layer using a two-sided label-permutation test on the difference of group means, correcting for multiple comparisons across layers using the false discovery rate (see Methods).

Across models, perceptual and semantic dimensions showed broadly similar emergence profiles, with prediction accuracy generally increasing across model depth (Figure 4A). However, perceptual dimensions were more strongly predictable than semantic dimensions in several simpler representations: this difference was already present in raw pixels and remained evident in the untrained AlexNet and feedforward CORnet, where several layers showed significant perceptual-semantic differences after FDR correction (Figure 4A). In contrast, this separation was reduced or absent in trained AlexNet, the recurrent and skip-connected CORnet models, and the transformer-based models, where the difference between perceptual and semantic profiles was not significant (Figure 4A). These results suggest that perceptual information carries a predictability advantage in stimulus statistics and simpler network representations, but that this distinction becomes less pronounced as representations are transformed by training and increasing model complexity.

**Figure 4.**
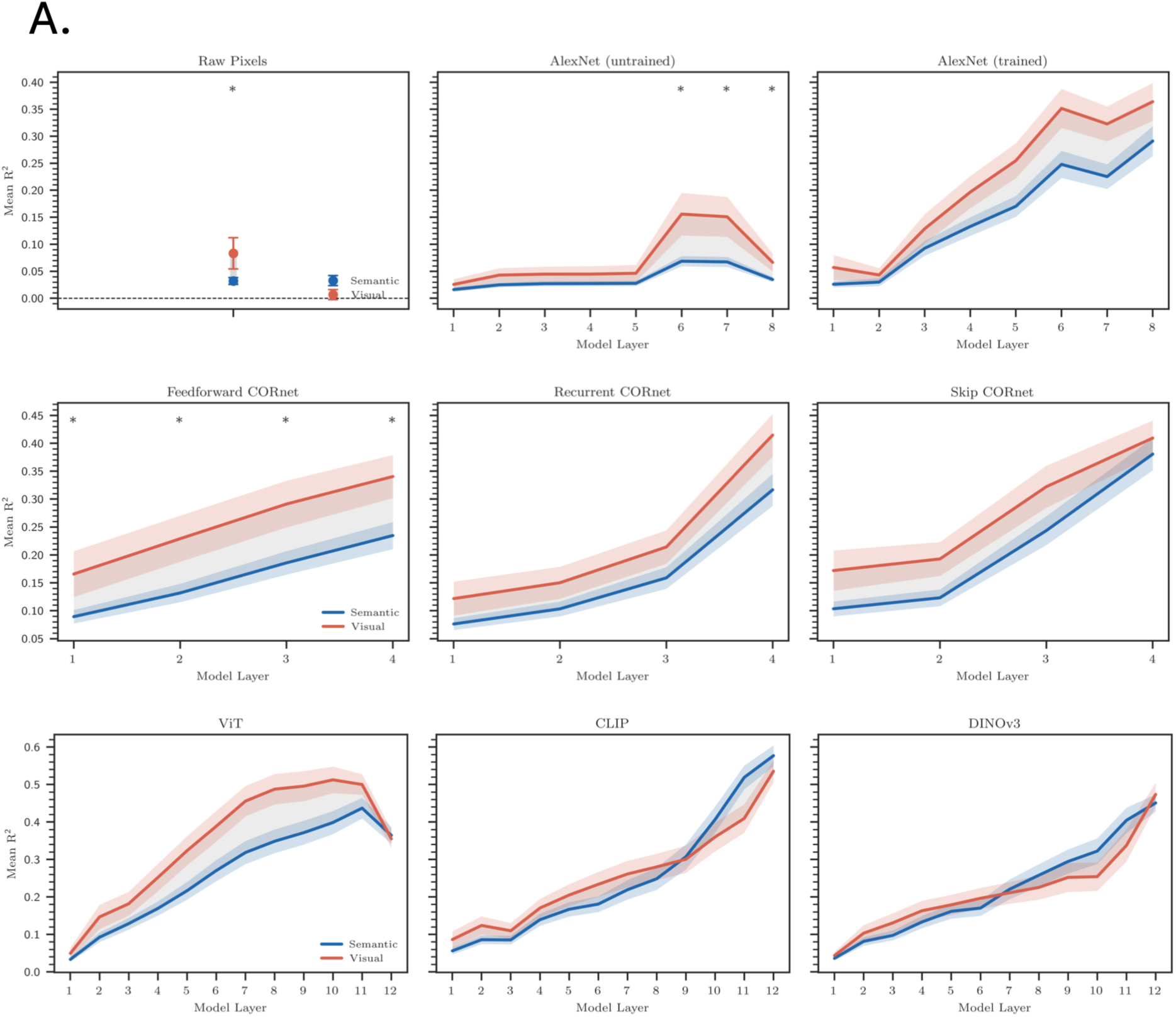
Perceptual-semantic differences are strongest in simpler representations. Mean prediction accuracy (R²) for semantic (blue) and perceptual (red) dimensions across raw pixels, untrained and trained AlexNet, CORnet architectures (feedforward, recurrent, and skip-connected), and transformer-based models (ViT, CLIP, DINOv3). Perceptual dimensions showed higher prediction accuracy than semantic dimensions in raw pixels, untrained AlexNet, and feedforward CORnet, with significant differences at several layers after FDR correction; this separation was reduced in trained AlexNet, recurrent and skip-connected CORnet models, and transformer-based models, where the difference between the two dimension types was not significant. Asterisks indicate significant perceptual-semantic differences after FDR correction (see Methods).

### Stimulus Statistics: Human-Derived Dimensions Generalise Across Domains

Having found systematic layer-wise prediction profiles for human-derived object dimensions, we next asked whether similar profiles are observed in a different perceptual domain: materials. Whereas object images are often structured by higher-level dimensions such as animacy and tool-relatedness, material images are more strongly associated with perceptual dimensions such as texture and geometry (Figure 5A). We therefore repeated our analyses on the STUFF material perception dataset (Schmidt et al., 2025), which focuses on natural and man-made materials (Figure 5A), allowing us to test whether layer-wise emergence profiles are specific to object-centric datasets or also observed for material-related dimensions.

**Figure 5.**
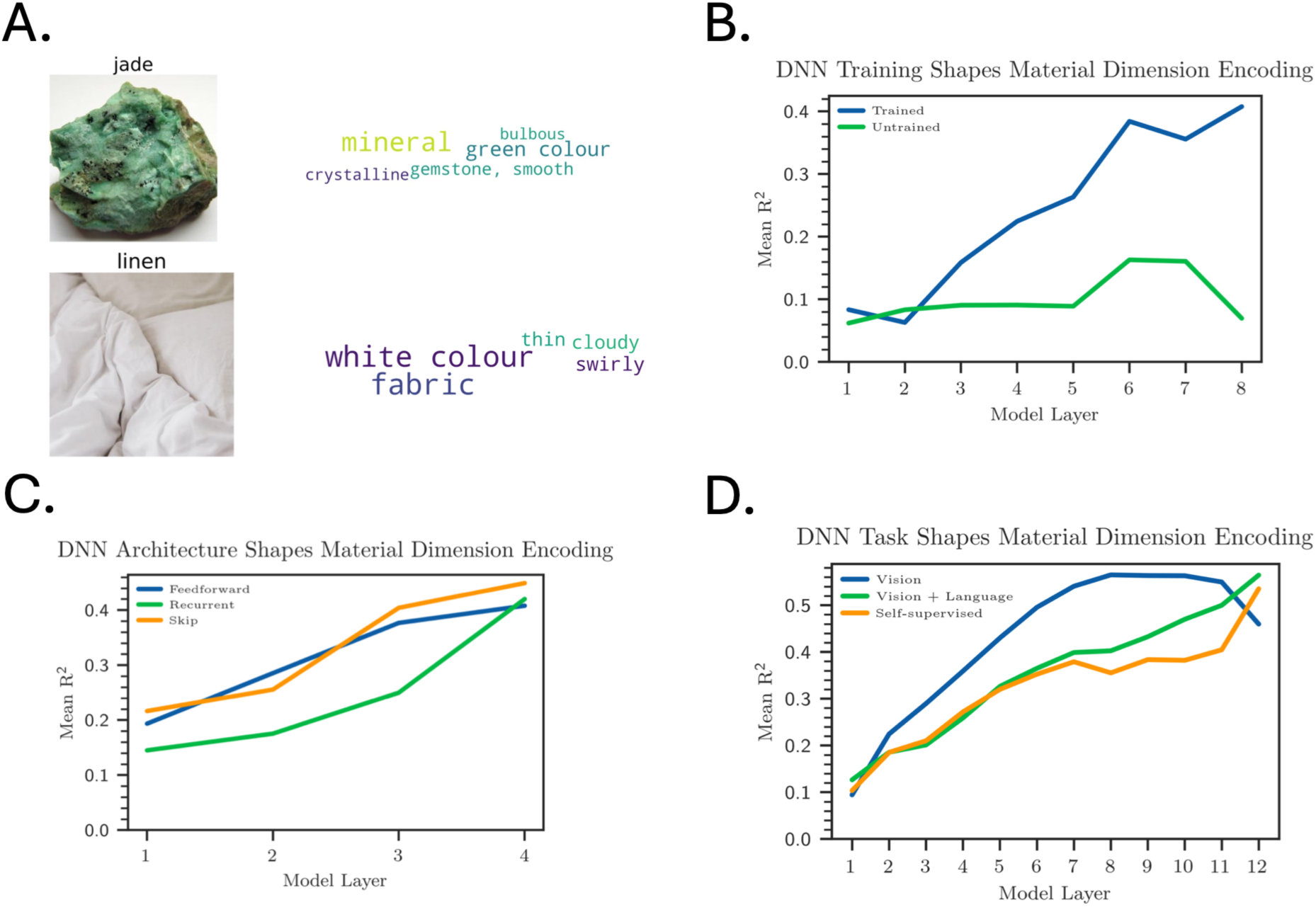
Predicting human-derived material dimensions across architectures and learning objectives. **A**. Example STUFF dataset images (jade, linen) with associated top-ranked human-derived material dimensions (e.g., mineral, crystalline, gemstone for jade; fabric, white colour, swirly for linen). Image credits: jade, Ra’ike/Wikimedia Commons (CC BY 3.0); linen, Sincerely Media/Unsplash. **B**. Mean prediction accuracy (R²) across AlexNet layers, comparing pretrained (blue) and untrained (green) models, showing that training substantially improves material dimension encoding. **C**. Comparison of network architectures (feedforward, recurrent, skip) on mean prediction accuracy (R²) across layers; the three architectures reach similar prediction in the final layer, and the recurrent model shows lower prediction than the other two in earlier layers. **D**. Comparison of learning objectives (vision-only, vision + language, self-supervised vision) for ViT-based models. All models show robust material dimension encoding, with vision-only slightly outperforming multimodal and self-supervised objectives in later layers.

Across models and architectures, human-derived material dimensions were robustly predictable from DNN activations and followed a clear layer-wise profile: prediction was weak in early layers, strengthened in mid-layers, and reached maximal prediction performance toward the top of the network (Figure 5B–D); relative to the THINGS object dataset, this profile was shifted toward earlier layers, though it still peaked toward the top of the network in absolute terms. These results suggest that emergence profiles are not limited to object dimensions, but can also be observed for human-derived material dimensions.

## Discussion

### Emergence profiles as a framework for model-brain comparison

In this study, we investigated whether DNN activations contain information predictive of human-derived dimensions in systematic layer-wise emergence profiles, and how architecture, learning objective, dimension type, and stimulus statistics shape these profiles across DNN layers. We found that trained networks produced clear emergence profiles, operationalised as trajectories of prediction performance across layers. Information predictive of human-derived dimensions was largely absent or weak in early layers, strengthened across intermediate layers, and often peaked in later layers. This layer-wise increase is a central finding in its own right, and it depends on training: it was absent in untrained AlexNet, which has the same depth as its trained counterpart, so depth alone does not produce it. Beyond this broad layer-wise pattern, the four factors we tested were systematically associated with differences in the shape of emergence profiles: recurrence and skip connections with greater strength and persistence of dimension prediction, learning objective with a different distribution of dimension-relevant information across layers, dimension type with a predictability advantage that was present in simpler representations but diminished in trained, more complex models, and stimulus statistics with differences in where human-derived dimensions were most strongly predictable. Among trained models, for example, prediction kept rising to the final layer in CLIP and DINffv3 but declined there in the supervised ViT, and recurrence and skip connections were associated with stronger prediction at different points in the network. Because prediction was resolved by layer and dimension, such differences can be located within the network, which a single-layer measure of decodability cannot do.

### Emergence profiles are not explained by architectural depth and dimensionality

Our findings suggest that systematic emergence profiles are not simply given by stimulus statistics or architectural depth alone, but depend strongly on network training. This is broadly consistent with human vision, where simple features appear earlier and more complex information builds up later (Carlson et al., 2013; DiCarlo et al., 2012; Robinson et al., 2023). Our results support a layer-wise increase in decodability, but not a mapping in which perceptual dimensions peak in earlier layers and semantic dimensions in later ones, since perceptual and semantic dimensions showed broadly similar trajectories in trained networks. Prediction from raw pixels revealed that some human-derived dimensions were already partly predictable from stimulus statistics. Untrained networks showed slightly stronger prediction for some dimensions, likely reflecting architectural biases, but still lacked clear progressive layer-wise prediction profiles. Only trained models produced clear emergence profiles in which information predictive of human-derived dimensions gradually strengthened across layers and peaked in later stages. Previous work has shown that DNNs contain information related to human-derived dimensions (Mahner et al., 2025) and broadly align with the human visual hierarchy (Cichy et al., 2016), but these studies largely examined dimension decodability at a single layer or as overall alignment, rather than tracking how it changes across layers. By introducing emergence profiles, we extend this work by showing not only that human-derived dimensions can be predicted from DNN activations, but also where in the network this prediction is strongest. This links findings from human vision (Contier et al., 2024; Teichmann et al., 2023) with layer-wise changes in artificial networks, while remaining agnostic about whether DNNs and humans share the same representational axes or geometry.

A related concern is whether prediction accuracy alone could be explained by a more generic account: later layers tend to carry higher-dimensional representations in general, so part of the layer-wise increase we observe could in principle reflect increasing representational complexity with depth rather than information organised specifically around human-derived dimensions. Our untrained-network control speaks against a purely architectural version of this account: untrained networks share the same depth and dimensionality at every layer as their trained counterparts (see Methods) yet did not produce a clear layer-wise emergence profile, indicating that increasing representational complexity with depth is not sufficient on its own to produce the pattern we observe in trained networks. This does not, however, rule out the related possibility that emergence profiles track general task performance rather than human-derived dimensions specifically, since untrained networks also perform poorly on the task; disentangling representational complexity, task performance, and dimension-specific organisation would require models that vary in task performance independently of depth and training, which we leave for future work.

### Recurrence and skip connections are associated with stronger dimension-relevant information

Comparing models that differed in specific architectural mechanisms showed that recurrence and skip connections were systematically associated with stronger and earlier-emerging dimension prediction, suggesting that these mechanisms may increase the availability of dimension-relevant information across layers. Because CORnet-RT and CORnet-S also achieve higher object-recognition performance than CORnet-Z, our results cannot fully rule out that part of this strengthening reflects improved task performance rather than recurrence or skip connections specifically. We note, however, that the two mechanisms were associated with qualitatively distinct signatures rather than a uniform rescaling of a single performance-related profile: recurrence was primarily associated with greater prediction magnitude in the final layer, whereas skip connections were associated with an earlier onset of strong prediction in the network. This pattern is consistent with the two mechanisms contributing in distinct ways rather than through a single common factor such as overall task performance, though fully disentangling architecture from task performance would require comparing models matched on task performance but differing in architecture.

### Learning objectives are associated with differences in information distribution across layers

Different learning objectives were associated with different profiles, with supervised vision–language and self-supervised models showing peak prediction in deeper layers, whereas vision-only showed trajectories that declined in the final layers. The pattern across learning objectives also differed between stimulus sets, in a way worth situating relative to prior work: material dimensions were robustly predicted by vision-only supervised models, whereas Storrs et al. (2021) argued that self-supervised learning is necessary to capture human-like material perception, particularly gloss. In our results, vision-only supervised training slightly outperformed self-supervised and vision–language objectives in predicting human-derived material dimensions in later layers. One possibility is that the broader set of material dimensions used here (spanning properties such as colour, texture, and surface appearance rather than gloss specifically) is more strongly tied to texture statistics that supervised classification objectives can also capture, without requiring the reconstruction-like pressure that favours self-supervised representations for finer-grained material properties such as gloss. Alternatively, supervision may simply not be as critical for material perception in general as proposed for gloss specifically.

### Perceptual versus semantic differences diminish in more complex models

Perceptual dimensions showed a predictability advantage over semantic dimensions in raw pixels, untrained AlexNet, and feedforward CORnet, but this advantage was reduced or absent in trained AlexNet, recurrent and skip-connected CORnet, and all three transformer models. This merits comment in light of Mahner et al. (2025), who found that DNN representational geometry is dominated by visual rather than semantic dimensions relative to humans. Our results are not straightforwardly in tension with this account: representational alignment (the extent to which a network’s internal geometry privileges certain dimensions) and decodability (whether a dimension’s value can be linearly recovered from a network’s activations) are distinct measures, and the pattern we observe describes when a perceptual advantage in decodability is present rather than whether a network’s geometry privileges visual dimensions overall. We are cautious about attributing the diminishing perceptual advantage specifically to training, since architecture and training are confounded across our model comparisons (e.g. the transformer models differ in learning objective, architecture details, and training data); we therefore describe this pattern as associated with more complex, trained representations rather than as a direct consequence of training itself. We also note that our tests may be underpowered to detect smaller differences, given the comparatively small number of dimensions classified as perceptual (15) relative to semantic (44); the absence of a significant separation in trained and more complex models should be interpreted as an absence of evidence for a difference of the size observed in simpler representations, rather than as evidence that no difference exists.

### Considerations and implications

Two further caveats apply across the factors we investigated. First, our results sample only a small fraction of the possible variation within each factor, and these findings should be interpreted as showing that each factor can be associated with differences in emergence profiles rather than as a complete account of how it does so. Second, dimensions vary substantially in how much variance they explain in the original human similarity judgments (Hebart et al., 2020): dimensions that explain more variance in human behaviour tend to have less noisy, more reliably estimated per-object weights, providing a cleaner prediction target for our regression models. Because prediction accuracy partly reflects how reliably a dimension’s per-object weights were estimated in the first place, dimensions with higher behaviourally explained variance may show stronger apparent emergence profiles for reasons unrelated to how that information is organised across DNN layers. These results indicate that emergence profiles are not fixed properties of a network, but layer-wise prediction trajectories that vary with design choices and stimulus statistics. By showing how information predictive of human-derived dimensions changes under different conditions, this approach clarifies factors associated with layer-wise prediction profiles in DNNs and provides hypotheses for how comparable factors may influence dimension-relevant information in the brain. Testing these predictions with neural data, such as fMRI or EEG, could reveal not only where models and the brain converge in their prediction profiles, but also where they differ in representational geometry.

In conclusion, this study shows that DNNs exhibit emergence profiles for human-derived dimensions, defined as layer-wise trajectories of prediction performance. These profiles are robust across the trained networks tested here and are systematically related to four factors: architecture, learning objective, dimension type, and stimulus statistics. Together, these findings show that information predictive of human-derived dimensions becomes available across DNN layers in structured ways, providing a layer-resolved framework for comparing artificial networks with broad progressions observed in human visual processing.

## Methods

All code for this work is available under https://github.com/Flo-Burger/Layer-Wise_Emergence_of_Human-Derived_Object_Dimensions_in_Deep_Neural_Networks. The THINGS dataset is available under https://things-initiative.org. To access the STUFF dimensions, please see https://osf.io/5gr73/.

### Datasets

We used two large-scale behavioural datasets providing human-derived dimensions. The THINGS database (Hebart et al., 2019, 2020) contains 26,107 images from 1,852 everyday object concepts. Human similarity judgments were modelled to derive 66 human-derived object dimensions that capture properties such as animacy, tool-relatedness, and category structure (S1 Table). Dimension weights were established only for the first exemplar of each concept, meaning that 1,852 images with associated dimensions were available for our analyses. In contrast, the STUFF database (Schmidt et al., 2025) contains 600 images from 200 natural and artificial material categories, such as stone, wood, and fabric. Using the same behavioural-modelling framework, 36 human-derived material dimensions were derived that primarily capture properties such as colour, texture, and surface appearance (S1 Table). In this dataset, human-derived material dimensions were established for all images, allowing us to include the entire set in our analyses. For both datasets, all images were resized to 224 × 224 pixels and normalized to the ImageNet mean and variance before being presented to the networks.

### Models

To examine how architectural mechanisms and learning objectives shape layer-wise emergence profiles for human-derived dimensions, we tested a range of deep neural network models differing in structure and learning paradigm. For each model, activations were extracted layer by layer, or block by block for transformer models, allowing us to track the trajectory of prediction performance for human-derived dimensions across model layers. All models were implemented in PyTorch (v1.13), with dimensionality reduction and regression carried out using scikit-learn (v1.4) and NumPy (v1.26). All analyses were run locally on a MacBook Pro 2024 with an M4 Pro chip.

As a baseline, we included raw image pixels as a control, testing whether human-derived dimensions could be predicted directly from stimulus statistics. We then analysed AlexNet (Krizhevsky et al., 2012), a widely studied convolutional network in cognitive science. By comparing AlexNet with random weights against its pretrained counterpart, trained on ImageNet-1k classification, we could test whether systematic layer-wise prediction profiles arise from architectural biases alone or depend on training.

To assess the influence of architectural mechanisms, we used the CORnet family of models (Kubilius et al., 2018), which implement a four-stage architecture inspired by the primate ventral visual hierarchy. CORnet-Z provides a purely feedforward baseline, CORnet-RT incorporates within-layer recurrence, and CORnet-S combines recurrence with within-layer skip connections. All three were used as the publicly released checkpoints, which were trained on the same dataset with the same objective (ImageNet-1k object classification; Kubilius et al., 2018). These models allowed us to test how recurrence and skip connections affect the strength and persistence of emergence profiles.

Finally, to investigate the role of learning objectives, we analysed three transformer-based models with comparable ViT-style architectures but different optimisation goals. We included ViT-Base (Dosovitskiy et al., 2021), trained with supervised image classification; the vision branch of CLIP (Radford et al., 2021), trained with an image–text contrastive objective; and DINOv3 (Siméoni et al., 2025), trained using a self-supervised objective. All three models were used as publicly released, pretrained checkpoints rather than models we trained or fine-tuned ourselves, and share the same core patch-embedding-plus-transformer-block architecture; they differ not only in learning objective but also in their specific training datasets (e.g., ImageNet-1k for ViT-Base, a web-scale image–text corpus for CLIP, and a large curated image corpus for DINOv3). Because learning objective and training data are confounded in these off-the-shelf models, differences between them should be interpreted as reflecting the combined influence of both factors rather than learning objective alone; disentangling them would require training matched models from scratch, which we leave for future work. For these models, we extracted the representation of the CLS (classification) token, a learned token prepended to the input sequence whose final-layer representation is conventionally used as a summary embedding of the whole image, from each transformer block. These models allowed us to test how different learning objectives influence the strength and distribution of layer-wise prediction profiles for human-derived dimensions.

### Feature Extraction and Dimension Prediction

For each dataset–model combination, we extracted activations from every layer. For convolutional neural networks, this included all convolutional and fully connected layers. For transformer models, this included the CLS token representation from each transformer block. These activations were reduced in dimensionality using principal component analysis (PCA) separately for each layer. For each layer, the PCA-transformed activations formed a feature matrix in which rows corresponded to images and columns corresponded to principal components, providing the input to the predictive model.

We tested different PCA configurations and regression models to ensure the robustness of results across analysis settings. To predict human-derived dimensions, we trained regression models separately for each dimension, with model performance quantified as the coefficient of determination (R²) averaged across 10-fold cross-validation. For each layer and each behavioural dimension, we fit a separate regression model, training on a subset of images and evaluating prediction accuracy on held-out images, rotating through folds such that each image served as test data once. This procedure yielded ten R² values for each dimension in each model layer, with each value corresponding to one cross-validation fold. These values were averaged to produce a single R² score for each layer–dimension pair, corresponding to one cell in the heat maps shown in the Results section. This procedure tested the ability of each layer’s activations to generalize to new stimuli when predicting a specific human-derived dimension. For our primary analyses, we selected the number of principal components separately for each layer as the minimum number needed to explain 80% of the variance in that layer’s activations, and used ridge regression (with regularisation strength selected via nested cross-validation) to predict each human-derived dimension. Allowing the number of retained components to vary by layer, together with ridge regularisation, lets the effective dimensionality of the predictor space adapt across layers while guarding against overfitting as the number of retained components grows relative to the number of images. We also examined a range of alternative configurations, including fixed component counts (50, 100) and a 95%-variance threshold, combined with unconstrained linear regression. These fixed-count linear-regression configurations were less stable, particularly at higher variance thresholds where the number of retained components approached or exceeded the number of available images, producing overfitting; results under ridge regression were stable across all tested PCA configurations (S2 Fig). Because several human-derived dimensions are conceptually nonnegative (e.g., an object cannot be meaningfully “negatively animal-related”), we additionally repeated this pipeline using nonnegative regression. This sensitivity analysis produced the same layer-wise pattern of results as the main ridge-regression analysis (S2 Fig). Throughout, we report R² values directly without additional normalization (e.g., min-max rescaling relative to the best-performing model or layer within an analysis), so that absolute magnitudes remain directly comparable across models and layers.

To summarize the layer-wise trajectory of each human-derived dimension, we defined its emergence profile as the sequence of R² values across layers. These profiles were then used to compare how architecture, learning objective, stimulus statistics, and dimension type shaped the layer-wise availability of information predictive of human-derived dimensions.

### Statistical Testing

We used permutation testing to determine whether prediction scores were above chance and whether differences between models were statistically reliable. First, for each model and layer, we built a null distribution of prediction accuracies by repeatedly shuffling the target dimension values 5,000 times and recomputing scores. This gave us a direct benchmark of what performance would look like under the null hypothesis, against which we compared the observed results. False discovery rate correction (FDR; Benjamini & Hochberg, 1995) was applied across all layers and human-derived dimensions to control for multiple comparisons.

In a second step, we used the same permutations to test differences between models. By pairing the permuted scores from two models, we generated a null distribution of the performance difference (ΔR²) for each human-derived dimension and layer. This allowed us to ask not only whether a given model performed above chance, but also whether prediction accuracy differed between two models for each human-derived dimension at each layer. These paired comparisons were again corrected for multiple comparisons using FDR. The same permutation-testing and FDR-correction procedure was applied to the linear- and ridge-regression panels of the supplementary robustness analysis (S2 Fig); nonnegative-regression panels beyond the original AlexNet/THINGS66d comparison report mean R² without permutation-based significance testing due to computational constraints.

## AI Use Section

We used Claude (Sonnet 5.5 and Opus 5.5; Anthropic) and ChatGPT (GPT-5.6 Sol; OpenAI) to assist with editing manuscript, response-letter text, and with writing analysis code. All AI-assisted text and code were reviewed, verified, and validated by the authors, who take full responsibility for the accuracy and content of this work.

## Acknowledgments

T. Grootswagers was supported by an Australian Research Council DECRA Fellowship (DE230100380), and M. Varlet was supported by an Australian Research Council Discovery Project grant (DP220103047). The funders had no role in study design, data collection and analysis, decision to publish, or preparation of the manuscript.

## Author Contributions

Conceptualization: Florian Burger, Tijl Grootswagers, Genevieve Quek, Manuel Varlet

Data curation: Florian Burger

Formal Analysis: Florian Burger

Software: Florian Burger

Supervision: Tijl Grootswagers, Genevieve Quek, Manuel Varlet

Methodology: Florian Burger, Tijl Grootswagers

Visualization: Florian Burger, Tijl Grootswagers, Genevieve Quek

Writing -original draft: Florian Burger

Writing -review & editing: Florian Burger, Tijl Grootswagers, Genevieve Quek, Manuel Varlet

## Supporting Information

S1 Table. Human-derived dimension labels and type classification for the 66 THINGS and 36 STUFF dimensions used in the heatmap figures.

S2 Fig. Robustness of dimension prediction across regression method and PCA configuration. Comparison of layer-wise prediction accuracy (R²) across three regression methods (unconstrained linear regression, ridge regression, and nonnegative regression) and two PCA configurations (80% variance threshold, 100 fixed components), for every model comparison (AlexNet, CORnet, and the three transformer models) on both datasets (THINGS66d, STUFF). Linear regression was less stable at high dimensionality, particularly on the smaller STUFF dataset, where the number of retained components approached or exceeded the number of available training images; nonnegative regression showed the same sample-size-driven instability at 100 fixed components on STUFF (e.g., mean R² ≈ −0.53 for AlexNet), while ridge regression remained stable across all tested configurations. Linear- and ridge-regression panels carry permutation-tested significance markers (FDR-corrected), as in the main figures; nonnegative-regression panels carry these markers only for the original AlexNet/THINGS66d comparison, and report mean R² without permutation-based significance testing elsewhere, to keep runtime tractable.

## References

Benjamini, Y., & Hochberg, Y. (1995). Controlling the False Discovery Rate: A Practical and Powerful Approach to Multiple Testing. Journal of the Royal Statistical Society Series B: Statistical Methodology, 57(1), 289–300. 10.1111/j.2517-6161.1995.tb02031.x

Bracci, S., Daniels, N., & Op De Beeck, H. (2017). Task Context Overrules Object- and Category-Related Representational Content in the Human Parietal Cortex. Cerebral Cortex, 27(1), 310–321. 10.1093/cercor/bhw419

Carlson, T., Tovar, D. A., Alink, A., & Kriegeskorte, N. (2013). Representational dynamics of object vision: The first 1000 ms. Journal of Vision, 13(10), 1–1. 10.1167/13.10.1

Cichy, R. M., Khosla, A., Pantazis, D., Torralba, A., & Oliva, A. (2016). Comparison of deep neural networks to spatio-temporal cortical dynamics of human visual object recognition reveals hierarchical correspondence. Scientific Reports, 6(1), 27755. 10.1038/srep27755

Contier, O., Baker, C. I., & Hebart, M. N. (2024). Distributed representations of behaviour-derived object dimensions in the human visual system. Nature Human Behaviour, 8(11), 2179–2193. 10.1038/s41562-024-01980-y

Conwell, C., Prince, J. S., Kay, K. N., Alvarez, G. A., & Konkle, T. (2024). A large-scale examination of inductive biases shaping high-level visual representation in brains and machines. Nature Communications, 15(1), 9383. 10.1038/s41467-024-53147-y

DiCarlo, J. J., Zoccolan, D., & Rust, N. C. (2012). How Does the Brain Solve Visual Object Recognition? Neuron, 73(3), 415–434. 10.1016/j.neuron.2012.01.010

Dosovitskiy, A., Beyer, L., Kolesnikov, A., Weissenborn, D., Zhai, X., Unterthiner, T., Dehghani, M., Minderer, M., Heigold, G., Gelly, S., Uszkoreit, J., & Houlsby, N. (2021). An Image is Worth 16x16 Words: Transformers for Image Recognition at Scale (arXiv:2010.11929). arXiv. 10.48550/arXiv.2010.11929

Groen, I. I., Greene, M. R., Baldassano, C., Fei-Fei, L., Beck, D. M., & Baker, C. I. (2018). Distinct contributions of functional and deep neural network features to representational similarity of scenes in human brain and behavior. eLife, 7, e32962. 10.7554/eLife.32962

Guclu, U., & Van Gerven, M. A. J. (2015). Deep Neural Networks Reveal a Gradient in the Complexity of Neural Representations across the Ventral Stream. Journal of Neuroscience, 35(27), 10005–10014. 10.1523/JNEUROSCI.5023-14.2015

Harel, A., Kravitz, D. J., & Baker, C. I. (2014). Task context impacts visual object processing differentially across the cortex. Proceedings of the National Academy of Sciences, 111(10). 10.1073/pnas.1312567111

Hebart, M. N., Dickter, A. H., Kidder, A., Kwok, W. Y., Corriveau, A., Van Wicklin, C., & Baker, C. I. (2019). THINGS: A database of 1,854 object concepts and more than 26,000 naturalistic object images. PLOS ONE, 14(10), e0223792. 10.1371/journal.pone.0223792

Hebart, M. N., Zheng, C. Y., Pereira, F., & Baker, C. I. (2020). Revealing the multidimensional mental representations of natural objects underlying human similarity judgements. Nature Human Behaviour, 4(11), 1173–1185. 10.1038/s41562-020-00951-3

Hong, H., Yamins, D. L. K., Majaj, N. J., & DiCarlo, J. J. (2016). Explicit information for category-orthogonal object properties increases along the ventral stream. Nature Neuroscience, 19(4), 613–622. 10.1038/nn.4247

Hubel, D. H., & Wiesel, T. N. (1962). Receptive fields, binocular interaction and functional architecture in the cat’s visual cortex. The Journal of Physiology, 160(1), 106–154. 10.1113/jphysiol.1962.sp006837

Kaniuth, P., Mahner, F. P., Perkuhn, J., & Hebart, M. N. (2024). A high-throughput approach for the eWicient prediction of perceived similarity of natural objects. bioRxiv. 10.1101/2024.06.28.601184

Kar, K., Kubilius, J., Schmidt, K., Issa, E. B., & DiCarlo, J. J. (2019). Evidence that recurrent circuits are critical to the ventral stream’s execution of core object recognition behavior. Nature Neuroscience, 22(6), 974–983. 10.1038/s41593-019-0392-5

Konkle, T., & Caramazza, A. (2013). Tripartite Organization of the Ventral Stream by Animacy and Object Size. Journal of Neuroscience, 33(25), 10235–10242. 10.1523/JNEUROSCI.0983-13.2013

Kriegeskorte, N. (2015). Deep Neural Networks: A New Framework for Modeling Biological Vision and Brain Information Processing. Annual Review of Vision Science, 1(1), 417–446. 10.1146/annurev-vision-082114-035447

Krizhevsky, A., Sutskever, I., & Hinton, G. E. (2012). ImageNet classification with deep convolutional neural networks. Communications of the ACM, 60(6), 84–90. 10.1145/3065386

Kubilius, J., Schrimpf, M., Nayebi, A., Bear, D., Yamins, D. L. K., & DiCarlo, J. J. (2018). CORnet: Modeling the Neural Mechanisms of Core Object Recognition. bioRxiv. 10.1101/408385

Mahner, F. P., Muttenthaler, L., Güçlü, U., & Hebart, M. N. (2025). Dimensions underlying the representational alignment of deep neural networks with humans. Nature Machine Intelligence, 7(6), 848–859. 10.1038/s42256-025-01041-7

Radford, A., Kim, J. W., Hallacy, C., Ramesh, A., Goh, G., Agarwal, S., Sastry, G., Askell, A., Mishkin, P., Clark, J., Krueger, G., & Sutskever, I. (2021). Learning Transferable Visual Models From Natural Language Supervision. In Proceedings of the 38th International Conference on Machine Learning (Vol. 139, pp. 8748–8763). PMLR. https://proceedings.mlr.press/v139/radford21a.html

Ren, Y., & Bashivan, P. (2024). How well do models of visual cortex generalize to out of distribution samples? PLOS Computational Biology, 20(5), e1011145. 10.1371/journal.pcbi.1011145

Robinson, A. K., Quek, G. L., & Carlson, T. A. (2023). Visual Representations: Insights from Neural Decoding. Annual Review of Vision Science, 9(1), 313–335. 10.1146/annurev-vision-100120-025301

Schmid, M. C., Mrowka, S. W., Turchi, J., Saunders, R. C., Wilke, M., Peters, A. J., Ye, F. Q., & Leopold, D. A. (2010). Blindsight depends on the lateral geniculate nucleus. Nature, 466(7304), 373–377. 10.1038/nature09179

Schmidt, F., Hebart, M. N., Schmid, A. C., & Fleming, R. W. (2025). Core dimensions of human material perception. Proceedings of the National Academy of Sciences, 122(10), e2417202122. 10.1073/pnas.2417202122

Schrimpf, M., Kubilius, J., Hong, H., Majaj, N. J., Rajalingham, R., Issa, E. B., Kar, K., Bashivan, P., Prescott-Roy, J., Geiger, F., Schmidt, K., Yamins, D. L. K., & DiCarlo, J. J. (2018). Brain-Score: Which Artificial Neural Network for Object Recognition is most Brain-Like? bioRxiv. 10.1101/407007

Siméoni, O., Vo, H. V., Seitzer, M., Baldassarre, F., Oquab, M., Jose, C., Khalidov, V., Szafraniec, M., Yi, S., Ramamonjisoa, M., Massa, F., Haziza, D., Wehrstedt, L., Wang, J., Darcet, T., Moutakanni, T., Sentana, L., Roberts, C., Vedaldi, A., … Bojanowski, P. (2025). *DINOv3* (arXiv:2508.10104). arXiv. 10.48550/arXiv.2508.10104

Spoerer, C. J., Kietzmann, T. C., Mehrer, J., Charest, I., & Kriegeskorte, N. (2020). Recurrent neural networks can explain flexible trading of speed and accuracy in biological vision. PLOS Computational Biology, 16(10), e1008215. 10.1371/journal.pcbi.1008215

Stoinski, L. M., Perkuhn, J., & Hebart, M. N. (2023). THINGSplus: New norms and metadata for the THINGS database of 1854 object concepts and 26,107 natural object images. Behavior Research Methods, 56(3), 1583–1603. 10.3758/s13428-023-02110-8

Storrs, K. R., Anderson, B. L., & Fleming, R. W. (2021). Unsupervised learning predicts human perception and misperception of gloss. Nature Human Behaviour, 5(10), 1402–1417. 10.1038/s41562-021-01097-6

Teichmann, L., Hebart, M. N., & Baker, C. I. (2023). Dynamic representation of multidimensional object properties in the human brain. bioRxiv. 10.1101/2023.09.08.556679

Thorat, S., Proklova, D., & Peelen, M. V. (2019). The nature of the animacy organization in human ventral temporal cortex. eLife, 8, e47142. 10.7554/eLife.47142

Yamins, D. L. K., Hong, H., Cadieu, C. F., Solomon, E. A., Seibert, D., & DiCarlo, J. J. (2014). Performance-optimized hierarchical models predict neural responses in higher visual cortex. Proceedings of the National Academy of Sciences, 111(23), 8619–8624. 10.1073/pnas.1403112111

Zhuang, C., Yan, S., Nayebi, A., Schrimpf, M., Frank, M. C., DiCarlo, J. J., & Yamins, D. L. K. (2021). Unsupervised neural network models of the ventral visual stream. Proceedings of the National Academy of Sciences, 118(3), e2014196118. 10.1073/pnas.2014196118

