## Supporting Information for "Layer-Wise Emergence of Human-Derived Object Dimensions in Deep Neural Networks"

### S1 Table. Human-derived dimension labels and type classification — THINGS (66 dimensions)

| Dimension | Label | Type |
|---|---|---|
| 1 | metallic/artificial | semantic |
| 2 | food-related | semantic |
| 3 | animal-related | semantic |
| 4 | textile | semantic |
| 5 | plant-related | semantic |
| 6 | house-related/furnishing-related | visual |
| 7 | valuable/precious | semantic |
| 8 | transportation-/movement-related | semantic |
| 9 | body-/people-related | semantic |
| 10 | wood-related/brown | visual |
| 11 | electronics/technology | semantic |
| 12 | colorful/playful | semantic |
| 13 | outdoors | semantic |
| 14 | circular/round | visual |
| 15 | paper-related/flat | semantic |
| 16 | sports-/playing-related | semantic |
| 17 | tools/handheld/elongated | semantic |
| 18 | fluid-related/drink-related | semantic |
| 19 | water-related | semantic |
| 20 | oriented/many things | unclear |
| 21 | powdery/earth-related | semantic |
| 22 | white | visual |

| Dimension | Label | Type |
|---|---|---|
| 23 | coarse pattern/many things | visual |
| 24 | red | visual |
| 25 | long/thin | visual |
| 26 | weapon-/danger-related | semantic |
| 27 | black | visual |
| 28 | household | semantic |
| 29 | feminine (stereotypical) | semantic |
| 30 | body part-related | semantic |
| 31 | tubular | visual |
| 32 | music-/hearing-/hobby-related/loud | semantic |
| 33 | grid-/grating-related | visual |
| 34 | repetitive/spiky | semantic |
| 35 | construction-/craftsmanship-related | mix visual-semantic |
| 36 | spherical/voluminous | visual |
| 37 | string-related/curved | semantic |
| 38 | seating-/standing-/lying-related | semantic |
| 39 | flying-/sky-related | semantic |
| 40 | bug-related/non-mammalian/disgusting | unclear |
| 41 | transparent/shiny/crystalline | mix visual-semantic |
| 42 | sand-colored | unclear |
| 43 | green | visual |
| 44 | bathroom-/wetness-related | semantic |

| Dimension | Label | Type |
|---|---|---|
| 45 | yellow | visual |
| 46 | heat-/fire-/light-related | visual |
| 47 | beams-/mesh-related | semantic |
| 48 | foot-/walking-related | semantic |
| 49 | box-related/container | semantic |
| 50 | stick-shaped/container | semantic |
| 51 | head-related | semantic |
| 52 | upright/elongated/volumous | semantic |
| 53 | pointed/spiky | visual |
| 54 | child-/toy-related/cute | unclear |
| 55 | farm-related/historical | semantic |
| 56 | seeing-related | semantic |
| 57 | medicine-/health-related | semantic |
| 58 | sweet / dessert-related | semantic |
| 59 | orange | semantic |
| 60 | thin/flat/wrapping | semantic |
| 61 | cylindrical/conical/cushioning | semantic |
| 62 | coldness-related/winter-related | semantic |
| 63 | measurement-related/numbers-related | semantic |
| 64 | fluffy/soft | semantic |
| 65 | masculine (stereotypical) | semantic |
| 66 | fine-grained pattern | unclear |

### S1 Table (cont.). Human-derived dimension labels — STUFF (36 dimensions)

| Dimension | Label |
|---|---|
| 1 | mineral |
| 2 | wood |
| 3 | metallic |
| 4 | fabric |
| 5 | white colour |
| 6 | grainy |
| 7 | crystalline |
| 8 | straw, fibrous |
| 9 | cloudy |
| 10 | small |
| 11 | beige colour, tan colour |
| 12 | viscous |
| 13 | black colour |
| 14 | brick |
| 15 | swirly |
| 16 | maroon colour |
| 17 | lines, long, tubular, vertical |
| 18 | mesh |

| Dimension | Label |
|---|---|
| 19 | yellow colour |
| 20 | multi-coloured |
| 21 | thin |
| 22 | bulbous |
| 23 | green colour |
| 24 | blue colour |
| 25 | shell |
| 26 | gemstone, smooth |
| 27 | sheet |
| 28 | hair |
| 29 | spongy |
| 30 | golden |
| 31 | red colour |
| 32 | round, smooth |
| 33 | cream colour |
| 34 | bumpy |
| 35 | hot |
| 36 | turquoise colour |

S2 Fig. Robustness across regression method and PCA setting — Alexnet

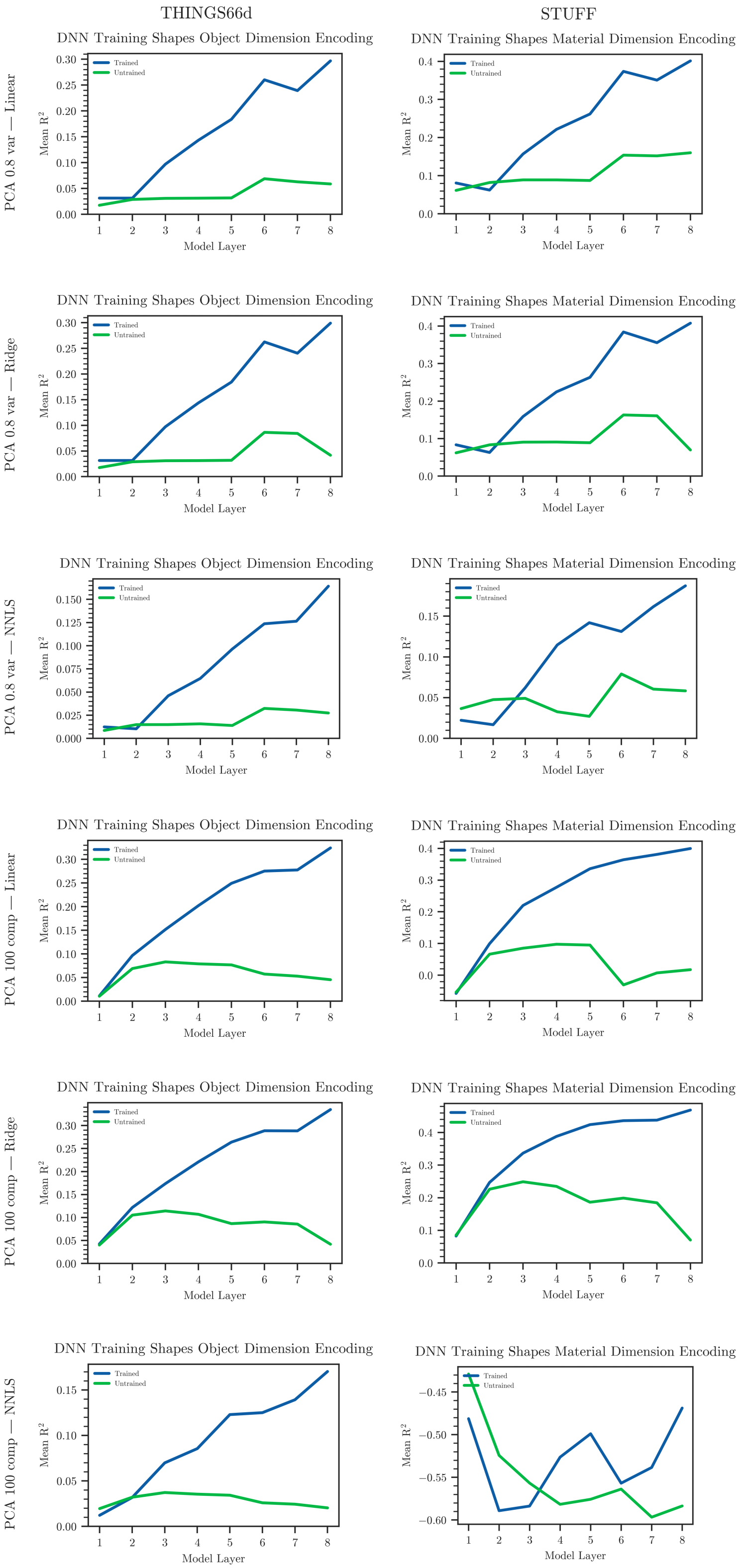


NNLS panels use no permutation-based significance testing except for AlexNet/THINGS66d; see Methods. NNLS, like unconstrained linear regression, becomes unstable at high PCA dimensionality on the smaller STUFF dataset (e.g. AlexNet, 100 components: mean $R^2 \approx -0.53$), consistent with the sample-size-driven overfitting mechanism already described in the Methods for linear regression.

S2 Fig. Robustness across regression method and PCA setting — Cornet

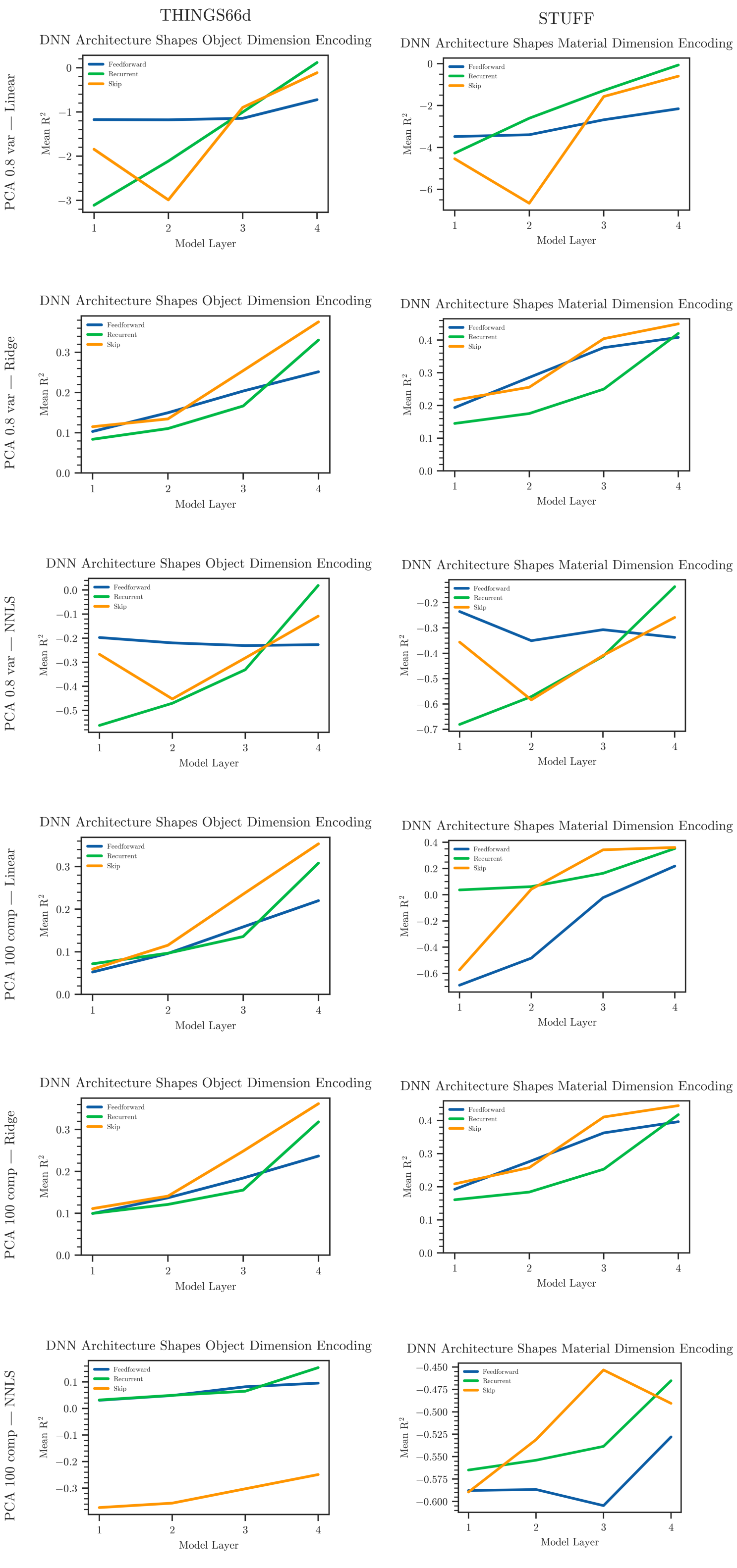


NNLS panels use no permutation-based significance testing except for AlexNet/THINGS66d; see Methods. NNLS, like unconstrained linear regression, becomes unstable at high PCA dimensionality on the smaller STUFF dataset (e.g. AlexNet, 100 components: mean $R^2 \approx -0.53$), consistent with the sample-size-driven overfitting mechanism already described in the Methods for linear regression.

S2 Fig. Robustness across regression method and PCA setting — Transformers

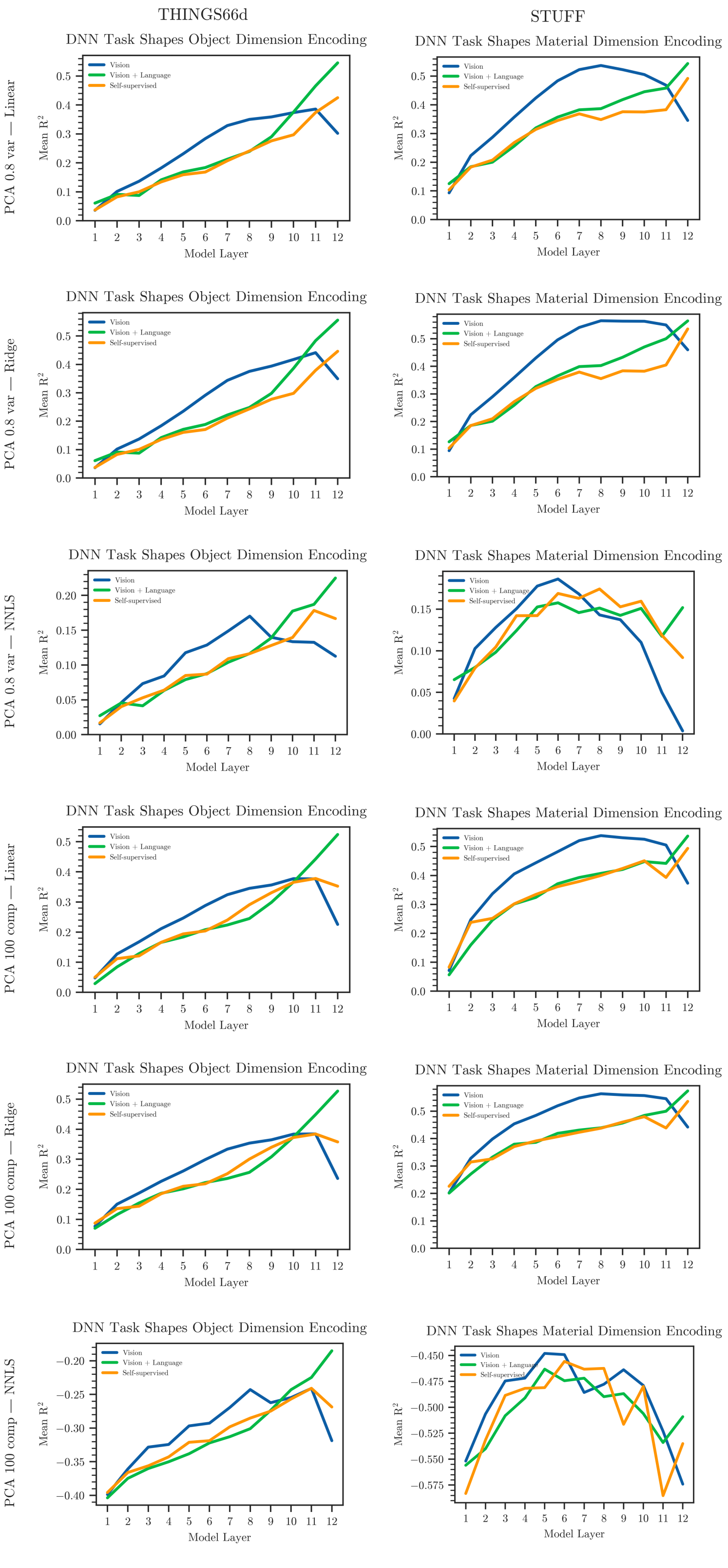


NNLS panels use no permutation-based significance testing except for AlexNet/THINGS66d; see Methods. NNLS, like unconstrained linear regression, becomes unstable at high PCA dimensionality on the smaller STUFF dataset (e.g. AlexNet, 100 components: mean $R^2 \approx -0.53$), consistent with the sample-size-driven overfitting mechanism already described in the Methods for linear regression.
